# Non-monotonic spatial structure of inter-neuronal correlations in prefrontal microcircuits

**DOI:** 10.1101/128249

**Authors:** Shervin Safavi, Abhilash Dwarakanath, Vishal Kapoor, Joachim Werner, Nicholas G. Hatsopoulos, Nikos K. Logothetis, Theofanis I. Panagiotaropoulos

## Abstract

Correlated fluctuations of single neuron discharges, on a mesoscopic scale, decrease as a function of lateral distance in early sensory cortices, reflecting a rapid spatial decay of lateral connection probability and excitation. However, spatial periodicities in horizontal connectivity and associational input as well as an enhanced probability of lateral excitatory connections in the association cortex could theoretically result in non-monotonic correlation structures. Here we show such a spatially non-monotonic correlation structure, characterized by significantly positive long-range correlations, in the inferior convexity of the macaque prefrontal cortex. This functional connectivity kernel was more pronounced during wakefulness than anesthesia and could be largely attributed to the spatial pattern of correlated variability between functionally similar neurons during structured visual stimulation. These results suggest that the spatial decay of lateral functional connectivity is not a common organizational principle of neocortical microcircuits. A non-monotonic correlation structure could reflect a critical topological feature of prefrontal microcircuits, facilitating their role in integrative processes.

**Significance statement:** The spatial structure of correlated activity of neurons in lower-order visual areas has been shown to linearly decrease as a measure of distance. The shape of correlated variability is a defining feature of cortical microcircuits as it constrains the computational power and diversity of a region. We show here for the first time a non-monotonic spatial structure of functional connectivity in the pre-frontal cortex where distal interactions are just as strong as proximal interactions during visual engagement of functionally similar PFC neurons. Such a nonmonotonic structure of functional connectivity could have far-reaching consequences in rethinking the nature and the role of prefrontal microcircuits in various cognitive states.

## Introduction

The intra-areal connectivity patterns of neural populations in the mammalian neocortex frequently repeat across cortical areas (1–4). Such canonical rules with general validity are important in understanding basic organizational principles and ensemble computations in cortical networks (1–3,5). Nevertheless, identifying deviations from these rules between sensory and higher order, association cortical areas could reveal properties leading to cortical network specialization and higher cognitive functions (1–3,5,6).

The spatial structure of intra-areal functional connectivity is frequently inferred by measuring the trial-by-trial correlated variability of neuronal discharges (spike count correlations) (7). One of the most well-established properties (a canonical feature) of intra-areal, mesoscopic, functional connectivity is a so called limited-range correlation structure, reflecting a monotonic decrease of spike count correlations as a function of spatial distance and tuning similarity (7–17). However, this distance-dependent decrease of correlations has been almost exclusively derived from recordings in primary sensory cortical areas or inferred from recordings in the PFC with various constraints like a rather limited scale (18) (see also Discussion section). As a result, it is currently unclear whether known differences in the structure of anatomical connectivity across the cortical hierarchy could also give rise to different spatial patterns of functional connectivity (19–22).

Specifically, the rapid spatial decay of correlations in sensory cortex is widely assumed to reflect a similar rapid decay in lateral anatomical connectivity and excitation (23). In early visual cortical areas, correlations rapidly decrease as a function of distance (12,14,17) (but also see (24,25)) in a manner that closely reflects anatomical findings about the limited spread and density of intrinsic lateral connections (19,26–29). However, lateral connections are significantly expanded in later stages of the cortical hierarchy, like the prefrontal cortex (PFC) (19,21,28–31). In this higher-order association area, lateral connections commonly extend to distances up to 7-8mm (28,29,31) while patches of connected populations are both larger and more distant from each other compared to sensory cortex (29,32). Although horizontal axons in macaque V1 can extend up to 4mm, they do not form clear patches, and for distances of 2-3mm laterally to the injection patch border, only a small number of cells are labelled in comparison to higher-order areas (19,27,29,33,34). In addition to the more extended intrinsic lateral connectivity, associational input from other cortical areas to the PFC also forms stripes with an average distance of 1.5mm and contributes to the spatial periodicities in lateral organization (35). Finally, the proportion of lateral excitatory connections is higher in the PFC (95%) compared to V1 (75%) (36).

Whether these significant differences in the structural architecture of the PFC compared to early sensory areas also result in a distinct spatial pattern of functional connectivity is currently unknown. Intuitively, higher probability of long-range lateral excitatory connections and stripe-like associational input patterns could give rise to strong spike count correlations across local and spatially remote populations with weaker correlations for populations in intermediate distances. To address this question, we recorded simultaneously the activity of large neural populations in the inferior convexity of the macaque PFC during both anesthetized and awake states using multi-electrode Utah arrays (37). In both anesthetized and awake states the spatial pattern of pairwise correlated variability was non-monotonic with significantly positive long-range correlations. A major source of non-monotonicity could be attributed to the spatial pattern of correlated variability between functionally similar neurons.

## Results

We used multi-electrode Utah arrays (4x4mm, 10x10 electrodes, inter-electrode distance 400μm, electrode length 1mm, Figure 1A) to record spiking activity from the inferior convexity of the ventrolateral prefrontal cortex (vlPFC) during repeated visual stimulation with movie clips in two anesthetized macaque monkeys (Figure 1B) and with sinusoidal gratings, drifting in 8 different directions, in two awake behaving macaques (Figure 1C). To evaluate the effect of structured visual input on correlated variability, we contrasted periods of visual stimulation to intertrial as well as spontaneous activity (long periods of neural activity without any task demands). Both anesthetized and awake state recordings resulted in the simultaneous monitoring of multiple, well isolated single units that remained stable for several hours of recording (Figure 1D). On average, in each dataset, we recorded from 103 ± 16 (mean ± S.E.M) single units and 5305 ± 1681 pairs during anesthesia (Figure S1A), and 107 ± 14 single units and 5758 ± 1675 pairs during wakefulness (Figure S1B).

**Figure 1.**
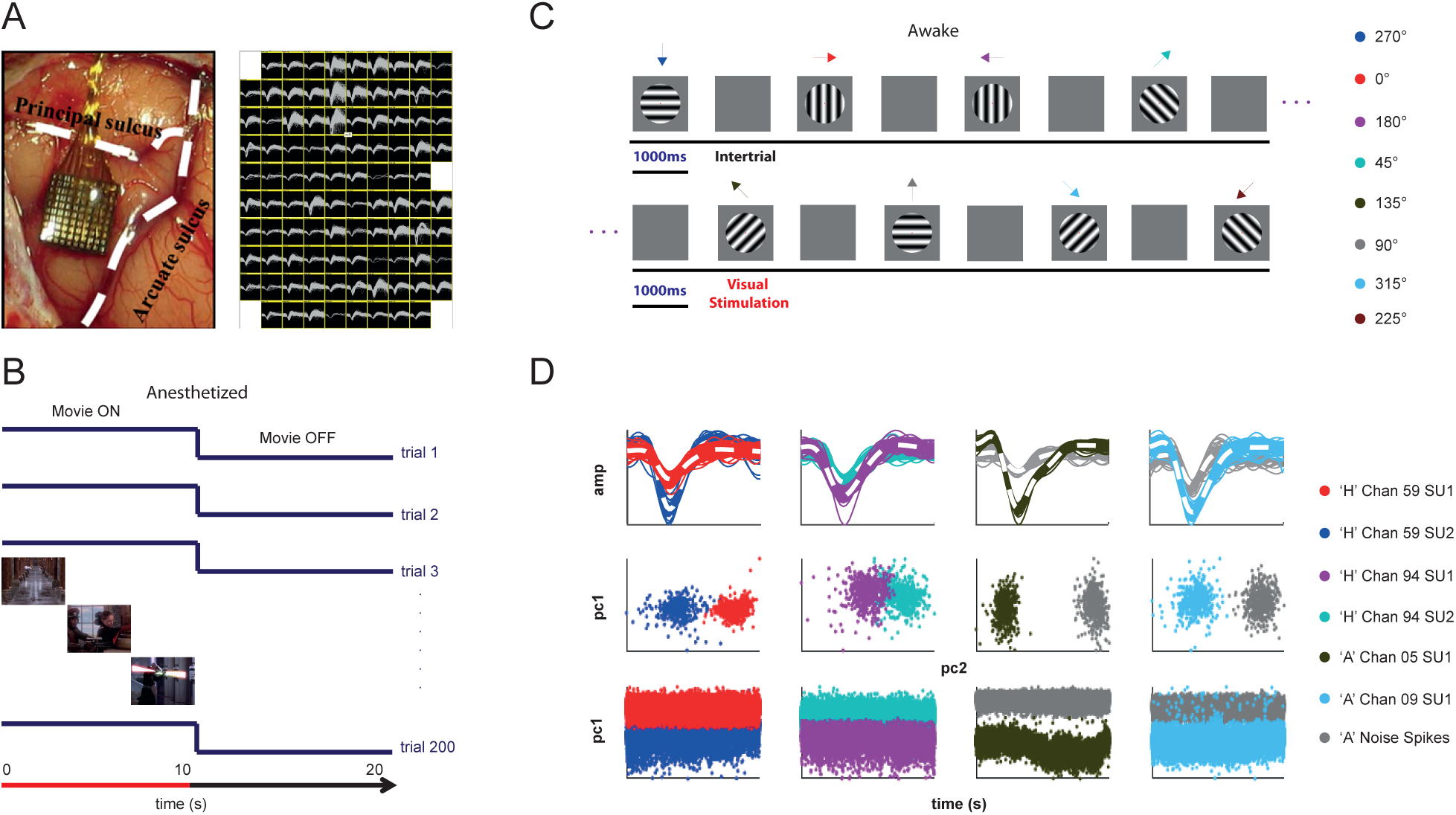
Implantation, visual stimulation and quality of single unit isolation. (A) Location of the implanted array with respect to arcuate and principal sulci and an example of typical waveforms acquired across the implanted cortical patch during a typical recording session in an awake animal. (B) Anesthetized visual stimulation protocol: 10 seconds of movie clip presentation was interleaved with 10 second long inter-trial (stimulus off) periods for 200 repetitions. (C) Awake visual stimulation protocol: The macaques initiated each trial by fixating on a red dot for 300ms, following which a drifting sinusoidal grating was presented monocular for 1000ms. After 1000ms of visual stimulation and a 300ms stimulus off period, liquid reward was delivered for successful fixation throughout the trial period. An intertrial period of 1000ms preceded the next trial. Each block of trials comprised eight different motion directions (exemplified by differently coloured arrows) presented in a random order. (D) Single unit isolation quality: Each column shows the activity recorded from 4 channels recorded in two different datasets, one from each of the two monkeys (monkey H and monkey A). 500 example waveforms for single units (shown as coloured clusters) and noise spikes (multi-unit activity shown as grey clusters) along with the mean waveform in dashed-white, and their corresponding first and second principal components (pc1 and pc2) are shown in the first and second row, respectively. In the last row the first principal component of all the waveforms in a cluster is plotted over time, demonstrating stability of recordings and single unit isolation for periods lasting ~3.5 to 4 hours.

### Spatial structure of correlated variability during anesthesia and wakefulness

It has been repeatedly shown that correlated variability of spike counts in early sensory, especially visual areas in different species decreases as a function of lateral distance, with strong interactions for proximal and progressively weaker interactions for distal (up to 4mm) neurons (14,15,17). We investigated the same relationship between spike count correlations (*r*_*sc*_) and lateral distance up to 4mm in the vlPFC.

Visual stimulation with movie clips during anesthesia or with drifting gratings during wakefulness, gave rise to a spatial pattern in the structure of correlated variability that was fundamentally different compared to early sensory areas: Strong and positive long-range ( > 2.5mm) correlations that were comparable to the average magnitude of local (up to 1mm) correlations and significantly weaker correlations for intermediate distances (red curves in Figure 2A and 2B and Figure S1A and S1B).

**Figure 2.**
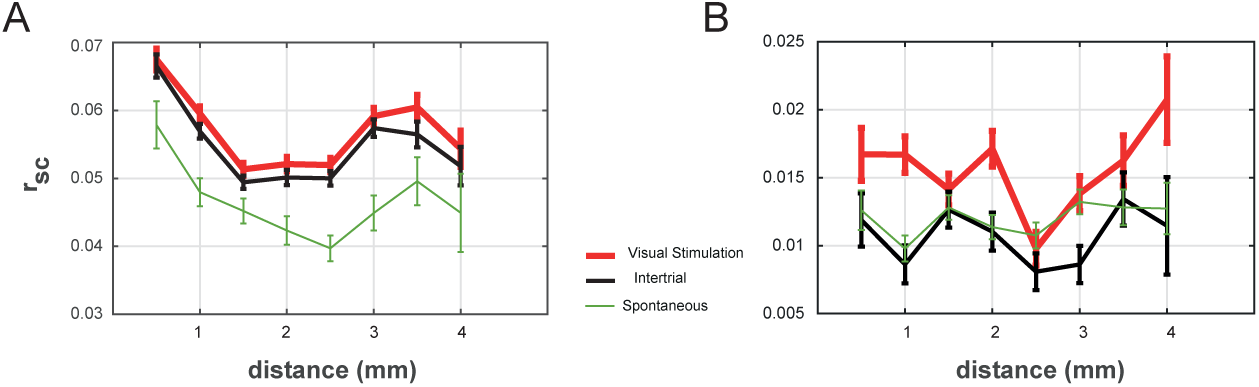
Spatial structure of correlated variability. (A) Spike count correlations (r_sc_) during visual stimulation (red), intertrial (black) and spontaneous activity (dark blue) as a function of lateral spatial distance (mm) between cell pairs for anesthetized state recordings (error bars represent mean ± SEM). (B) Same as (A) for awake state recordings.

We evaluated the statistical significance of non-monotonicity and long-range correlations by comparing the distributions of pairwise correlations in populations recorded from nearby (0.5mm for anesthetized and 1mm for awake), intermediate (2.5mm) and distant (3.5-4mm) sites during visual stimulation. The choice of these particular distance bins was based on the local extrema of correlated variability as a function of distance during visual stimulation. For all the comparisons made across these distance bins, in order to assess the significance of the differences in correlated variability, we used the Wilcoxon rank-sum test (unless otherwise mentioned explicitly). Moreover, as we made the comparisons across the three key distance bins, we assess the significance after a Bonferroni correction for multiple comparisons (corrected p-value = 0.0167). Summary statistics of Bonferroni corrected p-values are available in tables S1, S2 and S3 (for anesthetized and awake data respectively).

Average correlated variability between neurons located in intermediate distances was significantly lower compared to very proximal neurons in the anesthetized 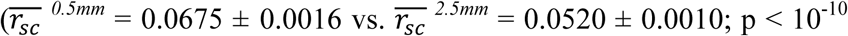, Figure 2A red curve and Figure S2A) state. In the awake recordings, correlations among nearby neurons, i.e. at a pairwise distance of 1mm showed a significant difference from those at the local minimum of the spatial correlation structure 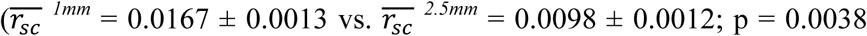, Figure 2B red curve and Figure S2D). Following this minimum, correlations during anesthesia significantly increased from 2.5mm to 3mm 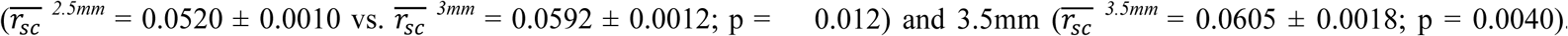. A similar increase in correlated variability for progressively more distant populations was also observed in the awake state, where correlations significantly increased from 2.5 to both 3.5mm 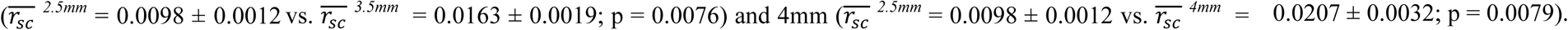 and 4mm.

In the awake state recordings, the average magnitude of correlations for distant populations, located 3.5-4mm apart, was not different from the respective magnitude for nearby pairs 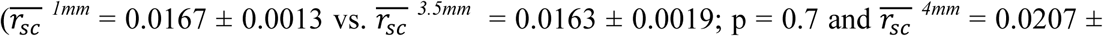 0.0032; p = 0.3, Figure 2B red curve and Figure S2D). In addition, both local and distant average correlations were significantly positive (p < 0.005, t-test). However, in the anesthetized recordings, despite the significant increase of correlations for distant neurons compared to intermediate distances, long-range correlations remained significantly lower compared to nearby neurons 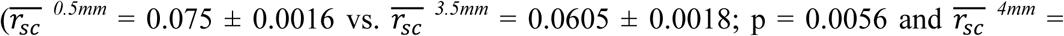 0.0544 ± 0.0027; p = 0.0008, Figure 2A red curve and Figure S2A), suggesting that anesthesia induced fluctuations had a non-homogeneous impact on the spatial structure of correlations. Such non-homogeneous, state-dependent weighting on the spatial structure of correlations has been reported in previous studies of primary visual cortex as well (12).

The non-monotonic structure in correlated variability could not be ascribed to random spatial variability in firing rates since it could be observed even when correlated variability was estimated for populations with matched geometric mean firing rates across lateral distances (Figure S3). Furthermore, to confirm that the intrinsic non-uniformity of spatial sampling with Utah arrays did not lead to the non-monotonic structure of correlated variability, we used a bootstrapping analysis of our spatial sampling (Figure S4). This analysis shows that equalized resampling of pairs across distance bins also results in a non-monotonic correlated variability structure (figure 2).

The decrease in correlations from nearby neuronal pairs (0.5mm in anesthetized state and 1mm in the awake state) to 2.5mm and the increase from 2.5 to 3.5 or 4mm was observed in both anesthetized and awake states. However, in the awake state recordings, we also observed an additional pronounced peak at 2mm (Figure 2B and Figure S2D). Lack of this peak at intermediate distances in our anesthetized recordings is compatible with other studies performed during anesthesia and provides further evidence for a non-homogeneous, state-dependent weighting on the spatial structure of correlated variability (12,14,17,25). These common features in the spatial structure of correlated activity across anesthetized 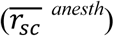 and awake states 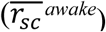 were observed despite significant differences in the average magnitude of correlations 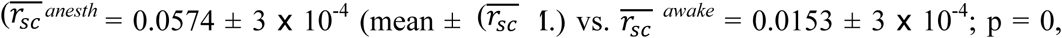 Figure 3A. Despite being very close to zero, average correlations during visual stimulation were significantly positive during the awake state (p < 10^−104^; t-test).

### Visual stimulation shapes the spatial structure of correlated variability

We evaluated the impact of structured visual stimulation on the spatial pattern of correlated variability by comparing correlations during visual stimulation with movie clips (during anesthesia) or drifting sinusoidal gratings (during wakefulness) to the respective pattern during intertrial and spontaneous activity periods. Compared to periods of intertrial activity visual stimulation resulted in a significant increase of correlated variability in both anesthetized 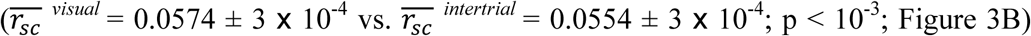 and awake recordings 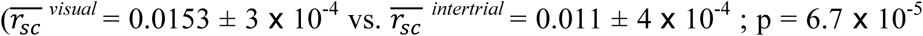, figure 3C.

**Figure 3.**
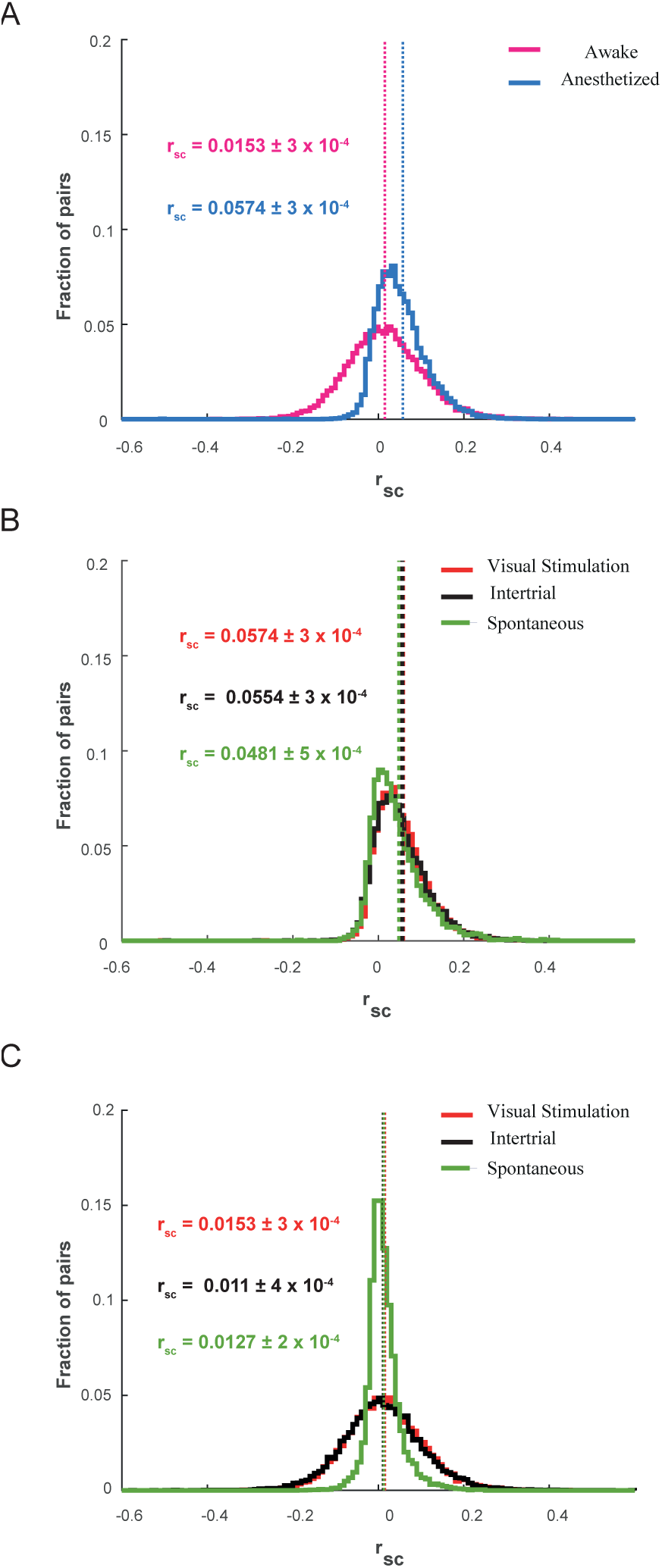
Distributions of correlated variability across different states and conditions. (A) Distribution of pairwise correlated variability (fraction of pairs) and mean values (dotted lines) during visual stimulation for anesthetized (blue) and awake (pink) recordings. Correlated variability was significantly stronger during anesthesia as a result of a shift in the distribution of pairwise correlations towards positive values. (B) Same as (A) for anesthetized state recordings during visual stimulation (red), intertrial (black) and spontaneous activity (green) periods. (C) Same as (B) for awake state recordings

Visual stimulation in the awake state significantly shaped a spatially inhomogeneous, non-monotonic structure of correlated variability. In striking contrast to the significant differences observed for the same lateral distances during visual stimulation, we found that correlations during the intertrial period were not different between local and intermediate 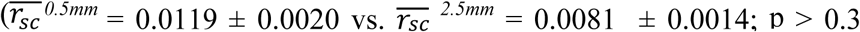 Figure 2B black curve and Figure S2B) or intermediate and distant populations 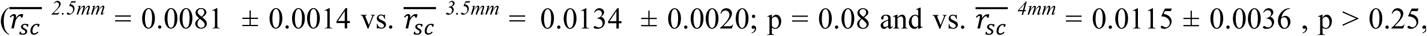, Figure 2B black curve and Figure S2B). Spatially homogeneous correlations were also observed during periods without any structured visual input or task engagement, in data collected during spontaneous activity (Figure 2B green curve). In these epochs, we also found no difference between local and intermediate correlations 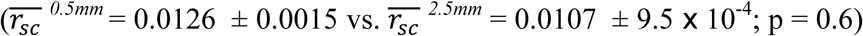 and very similar correlations between intermediate and distant populations 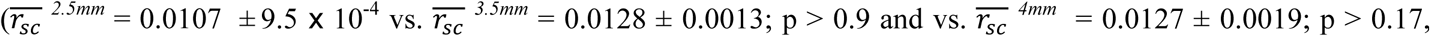, Figure 2B green curve).

We quantified the magnitude of spatial inhomogeneity in the structure of correlations across different conditions and states (see Experimental Procedures). A clear difference in the rate of changes in correlated variability was observed in awake state recordings (Figure 4A), where visual stimulation resulted in the strongest spatial variability and intertrial activity in the weakest (almost constant average correlation as a function of lateral distance). A similar spatial variability was also observed under anesthesia (Figure 4B); however, the average rate of change was comparable across the two conditions of visual stimulation and intertrial, but different during spontaneous activity. The difference in the structure of functional connectivity between visual stimulation and intertrial periods across anesthesia and awake states could be attributed to the lack of saccadic eye movements in intertrial periods during anesthesia. Saccadic eye movements reset visual perception (38) and their absence could create a persistent network state, showing no reset, resulting in very similar pattern of correlations during visual stimulation and intertrial periods.

**Figure 4.**
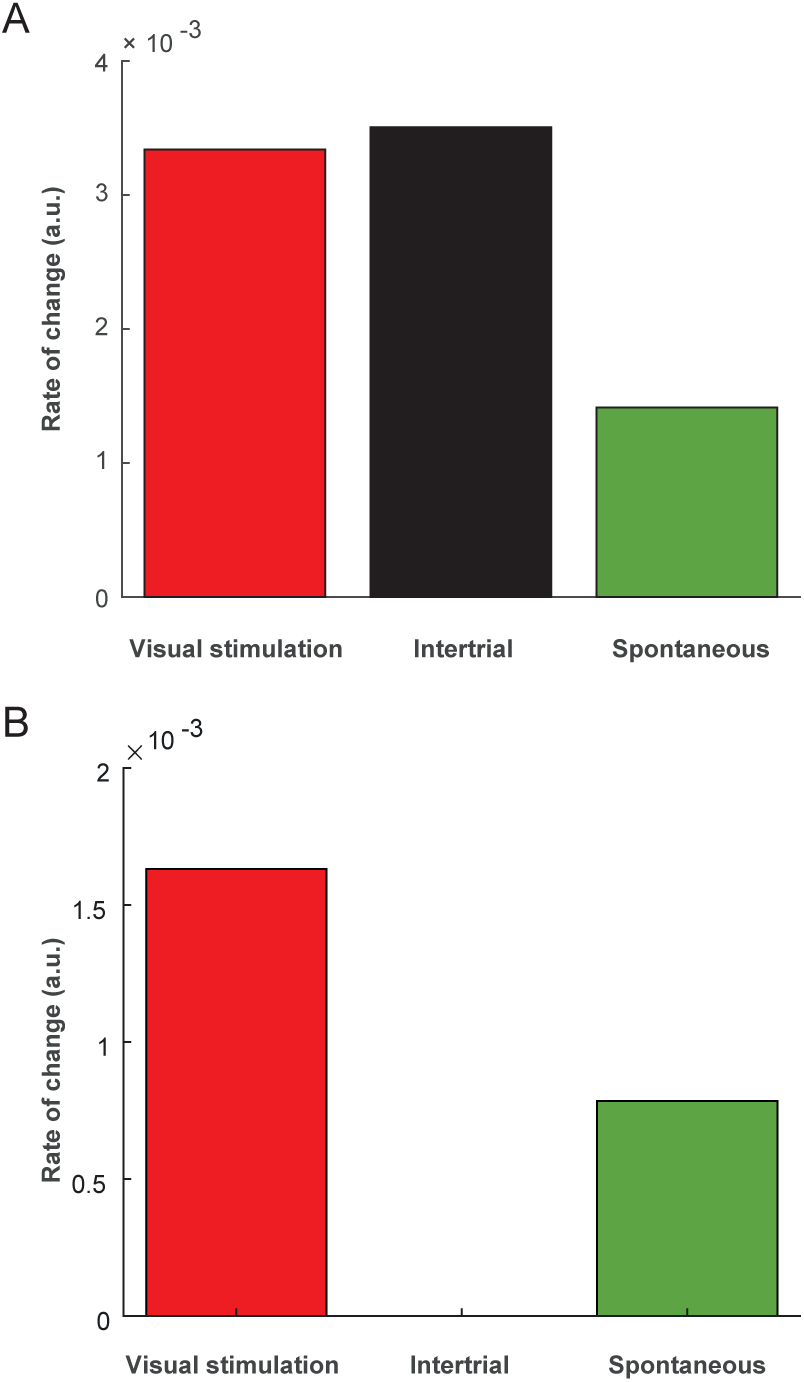
Quantification of spatial inhomogeneity in the structure of correlated variability. (A) Spatial inhomogeneity in the structure of correlations across different conditions during awake state recordings. Spatial inhomogeneity was quantified by computing the average of the absolute rate of change in the correlation structure across successive distance bins (only those rates significantly different in successive distance bins) (see also Experimental Procedures). (B) Same as (A) for anesthetized state recordings.

These results suggest that the spatial structure of correlated variability in PFC is inhomogeneous. The magnitude of inhomogeneity depended not only on the variation of global states such as wakefulness or anesthesia but most importantly on behavioural demands i.e. visual stimulation, intertrial (anticipation of the succeeding trial) or spontaneous activity (no behavioural load). Although traces of inhomogeneity in the spatial structure of correlations were observed during spontaneous activity or intertrial periods, structured visual stimulation during the awake state appeared to result in the strongest spatial inhomogeneity in the correlation structure.

### Prevalence of non-monotonic spatial structure in functionally similar populations

Lateral connectivity in PFC has been hypothesized to preferentially target neurons with functional similarities (e.g. similar spatial tuning), similar to iso-orientation columns in the visual cortex (Goldman-Rakic, 1995). Therefore, we next examined whether the source of the nonmonotonic correlated variability could be traced to populations of neurons that were modulated similarly by visual input. First, tuning functions for each recorded unit were obtained based upon the discharge response to sinusoidal gratings drifting in eight different directions (Figure 5). The correlation between tuning functions (signal correlation, *r*_*signal*_), provided a measure of functional similarity among the recorded pairs (see Experimental Procedures for more details). We analysed the relationship between the spatial structure of functional connectivity and functional similarity of pairwise responses (i.e. signal correlations). The relationship between signal correlations, noise correlations and inter-neuronal distance (Fig 6A) points to a stronger non-monotonic trend for pairs with positive signal correlations.

**Figure 5.**
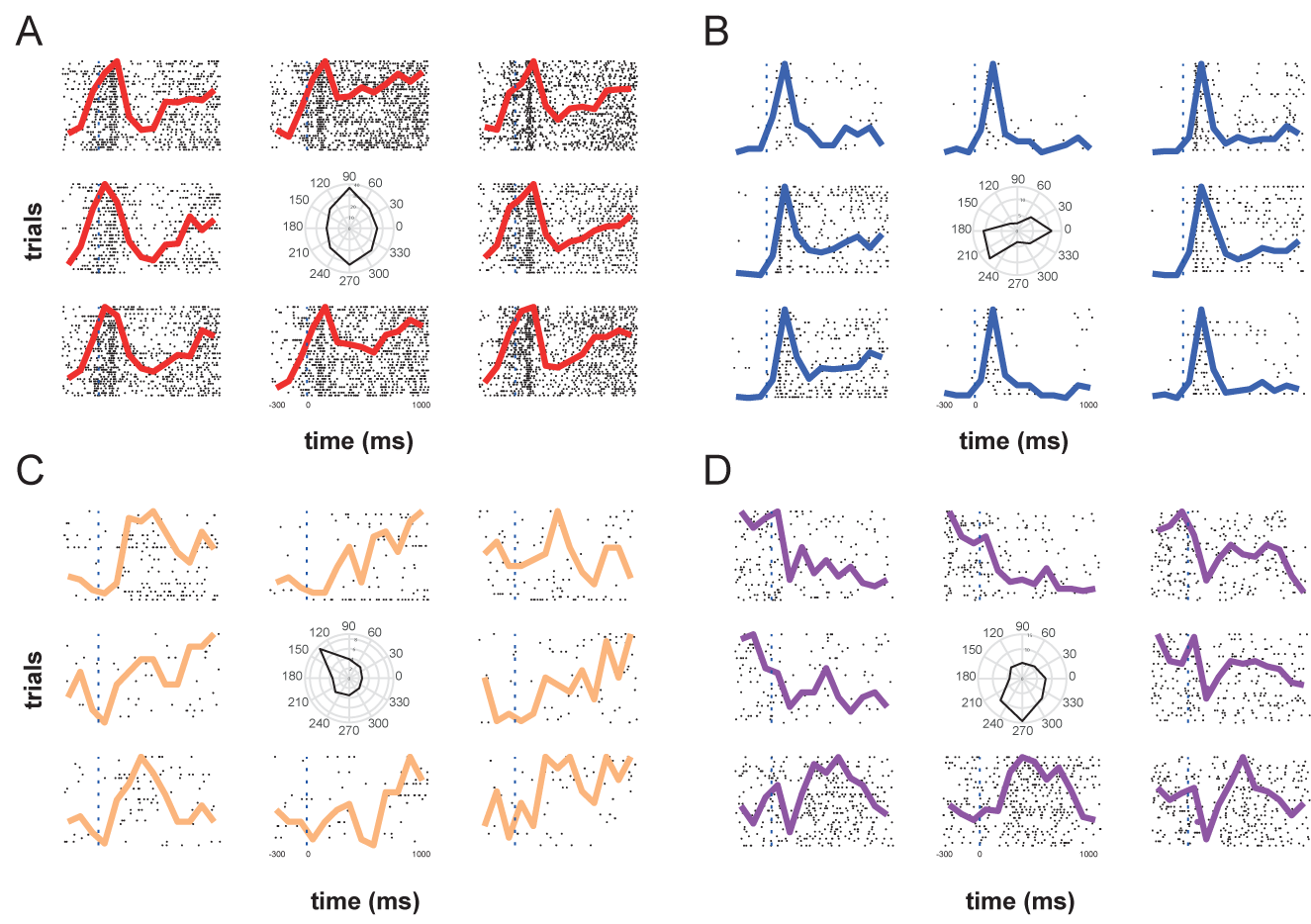
Visual modulation of single unit activity during wakefulness. Spike raster plots and overlaid peristimulus time histograms (PSTHs) for four single units (A-D) across all 8 orientations (0° - 315°). Polar plots for each unit show the preferred direction(s) of motion. The green lines in the centre indicate the resultant length and direction (see Supplemental Experimental Procedures, “Quantification of direction selectivity” for more details). The upper two panels are typical examples of bimodally tuned prefrontal units with significant responses for opposite directions of motion (orientation selective responses). The lower two panels are examples of sharper, unimodal responses for a particular direction of motion.

**Figure 6.**
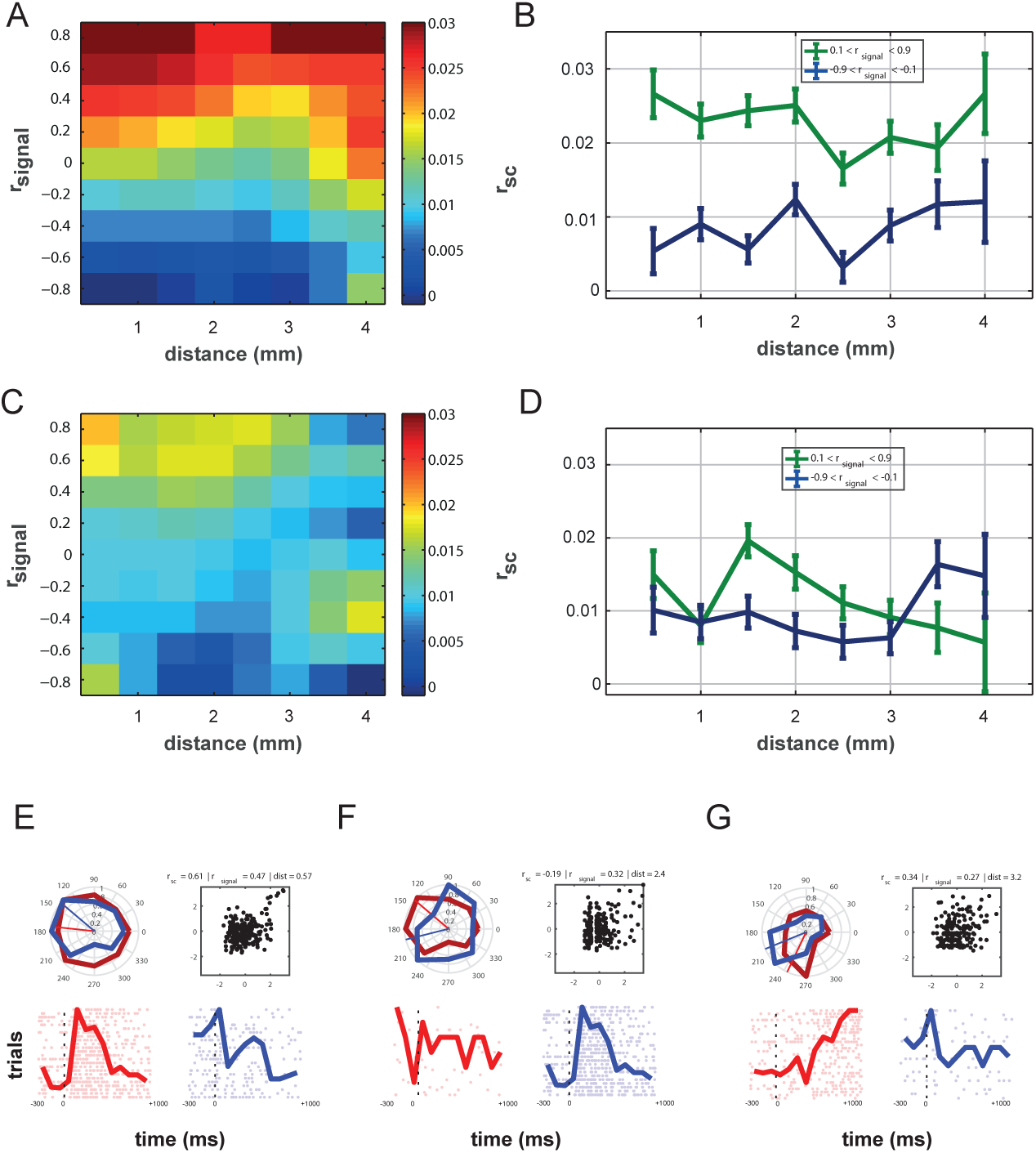
Effect of functional similarity on the spatial structure of correlated variability. (A) Correlated variability as a function of distance and signal correlation for the pooled data recorded during visual stimulation in the awake state. The colour of each pixel indicates the average correlated variability for pairs that their signal correlation and distance landed in the specific bin. Pixels containing less than 10 pairs are removed (white pixels). Correlated variability values are indicated by the colourbar at the right of the panel. Data were smoothed with a two-dimensional Gaussian (SD of 1 bin) for display purposes. (B) Correlated variability as a function of distance (similar to Figure 2A and B) among neuronal pairs with signal correlation higher than 0.1 and less than 0.9, i.e. the non-zero upper part of matrix represented in (A) with green line; and among pairs with signal correlation higher than −0.9 and less than −0.1 i.e. the non-zero lower part of matrix represented in (A) with blue line. (Mean ± SEM as error bars) (C-D) Same as (A) and (B) for the intertrial period. The signal correlation matrix is computed from the visual stimulation period and the correlated variability of these populations in the intertrial period is mapped onto the pixels in C. (E-G) Three example pairs with high signal correlations and high, low and high correlated variability from the nearest, the intermediate and the farthest distance bins, respectively. The polar plot shows the vector sum of the tuning for each neuron in a given pair while the scatter plots depict their z-score normalized responses. Example raster plots are overlaid with the peristimulus time histograms (PSTH) for the preferred direction of motion. Despite the sparseness in firing for some of the neurons, sharp tunings can be observed (compare raster plots with polar plots).

Specifically, we computed the noise correlation across distance bins for pairs with positive signal correlations (0.1 < *r*_*signal*_ < 0.9) during visual stimulation (Figure 6A,B). A nonmonotonic trend could be observed, however the differences between the first local maximum to the local minimum, and the local minimum to the next local maximum were marginally significant 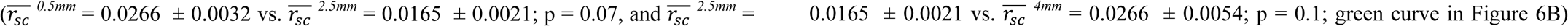. The nonmonotonic trend was also confirmed from fitting first and second-degree polynomials to these data. The adjusted-R^2^ goodness-of-fit measure for a line (first degree polynomial, monotonic) was −0.15 whereas the same measure for a quadratic function (second degree polynomial, the simplest non-monotonic function)(39) yielded a value of 0.3 pointing to the quadratic curve being a much better fit to the data - Figure 7A). Progressively higher thresholds for signal correlation, resulting in sampling populations with stronger functional similarity, did not qualitatively change these effects that were characterized by a significant decrease in intermediate distance (~2.5mm) correlations (Figure S5B,E,H).

**Figure 7.**
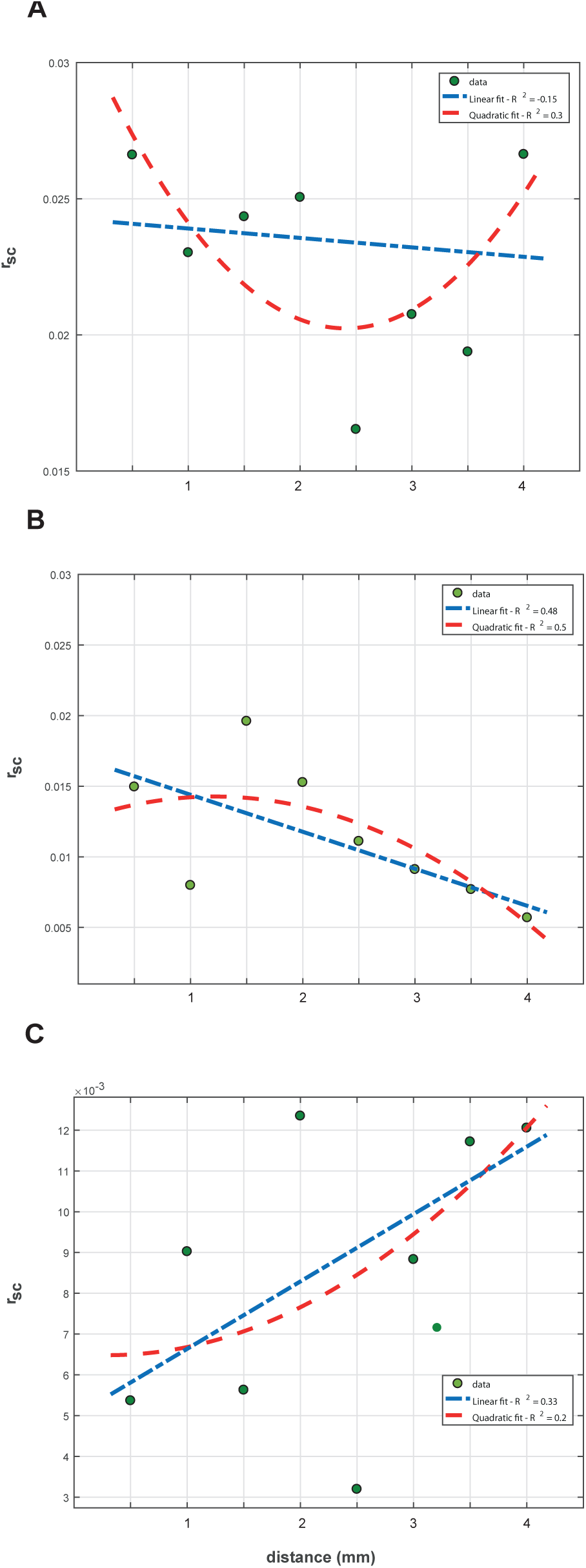
Fitting trends to the relationship between noise correlations and distance for functionally similar populations. (A) Linear (y = ax + b) and quadratic (y = ax^2^ + bx + c) trends fit to noise correlations as a function of distance for functionally similar neurons (0.1 < r_*signal*_ < 0.9) during visual stimulation periods. A negative adjusted R^2^ value for the linear fit and a positive R^2^ value of 0.3 for the quadratic fit clearly demonstrates a non-monotonic trend being shaped by visual stimulation. A positive symmetric convexity also points towards a strengthening of local and long-range connectivity during visual stimulation. (B) Linear (y = ax + b) and quadratic (y = ax^2^ + bx + c) trends fit to noise correlations as a function of distance for functionally similar neurons (0.1 < r_*signal*_ < 0.9) during intertrial periods. Very similar R^2^ values for both the fits (linear = 0.48 and quadratic = 0.5) demonstrates that the quadratic trend is not much different from a linearly decreasing trend. Moreover, the asymmetric negative convexity of the quadratic curve points to a lack of strengthening of local and long-range correlations when no visual stimulation is present. (C) Linear (y = ax + b) and quadratic (y = ax^2^ + bx + c) trends fit to noise correlations as a function of distance for functionally dissimilar neurons (-0.9 < r_*signal*_ < −0.1) during visual stimulation periods. The fitting results display a monotonically increasing trend of noise correlations as a function of distance. These pairs of neurons do not display the characteristic positive convexity shown by functionally similar neurons where local and distant populations have equivalent correlated variability, pointing to a different mechanism of functional connectivity driving this relationship

During intertrial periods, correlated variability of the same population of functionally similar neurons (functional similarity estimated during the visual stimulation period) was homogenous (Figure 6C,D). For positive signal correlations (0.1 < *r*_*signal*_ < 0.9) during visual stimulation, the strength of correlated variability between nearby and intermediate neurons was not significantly different 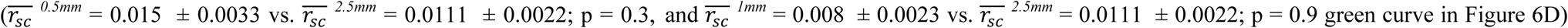 and correlations between neurons in intermediate and distant locations were also very similar 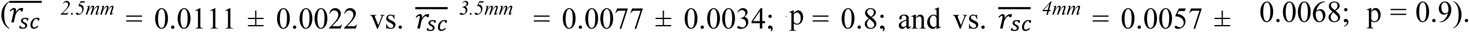. A similar fitting procedure as used for the data in the visual stimulation period was also employed to test for the observed trends in the inter-trial period. Fitting a line yielded an adjusted R^2^ value of 0.48 whereas fitting a quadratic function yielded an adjusted R^2^ value of 0.5 pointing to both fits being quantitatively similar (Fig. 7B). However, whereas the quadratic fit in the visual stimulation period yielded a U-shape curve that clearly displayed the equivalence between local and distant populations, this equivalence was not seen during the intertrial period where distant populations were weakly correlated compared to local populations.

Furthermore, when local (0.5-1mm) and distant (3.5-4mm) populations were pooled at a spatial resolution of 1mm, a clear and specific strengthening of correlated variability at the flanks was observed during structured visual stimulation epochs 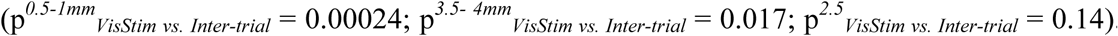. Taken together, the non-monotonic spatial structure of functionally similar populations during visual stimulation was stronger as compared to the more homogeneous structure (Fig. 7B) of the same population during the inter-trial period, pointing to a role of structured visual input in shaping the non-monotonic structure of correlated variability in functionally similar populations.

When the same fitting procedure as above was performed on pairs of functionally dissimilar neurons (i.e. −0.9 < r _*signal*_ < −0.1), a linearly increasing trend provided a slightly better fit as compared to a quadratic fit (adjusted R^2^ linear = 0.33, adjusted R^2^ quadratic = 0.2). However, when local and distant populations were binned as for the high signal correlation pairs, correlated variability in the flanks during visual stimulation and intertrial was identical - 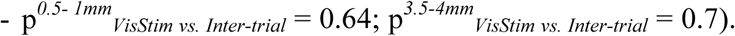. Examples of pairwise neuronal responses from neurons with similar signal correlations and sampled from short, intermediate and long lateral distances are presented in Figure 6E-G.

Several factors changed between the anesthetized versus awake animal experiments. For instance, recordings were performed in different monkeys, using different stimuli (movie clips versus moving grating), different data acquisition systems and spike extraction methods. Despite these differences, inter-neuronal correlations showed a similar spatial structure in both anesthetized and awake recordings with visual stimulation. More specifically, these results suggest that spatial inhomogeneities in the functional architecture of the PFC arise from strong local and long-range lateral functional interactions between functionally similar neurons, which are particularly pronounced during structured visual stimulation in the awake state.

## Discussion

### Spatial structure of prefrontal correlated variability and relationship to anatomical structure

Spatial decay in the strength of spike count correlations on a mesoscopic scale, up to 4 mm of lateral distance, is largely considered a canonical feature of functional connectivity. Our results suggest that this spatial decay is not observed in the PFC since nearby and distant neurons are correlated to the same degree, thus reflecting a fundamentally different lateral functional connectivity structure compared to primary sensory areas like V1 (14,17,24,25). Such a functional connectivity pattern is likely to directly reflect the underlying anatomical organization of prefrontal neural populations into spatially distributed clusters connected through local and long-range excitatory collaterals (28,32,36). Indeed, in awake state recordings, the spatially inhomogeneous correlation pattern reflected bumps of ~ 1.5-2 mm width (Figure 2B) which closely matches the spatial distribution (~1.5mm maximum width) of laterally labelled stripes of neuronal assemblies in supragranular prefrontal layers (Kritzer & Goldman-Rakic 1995; Pucak et al. 1996).

Although purely anatomical methods cannot identify functional similarities across connected populations (and vice versa), an influential hypothesis of structural connectivity in the PFC assumes that long-range excitatory collaterals target clusters of neurons with similar functional preference, like spatial tuning (40). Our results provide experimental evidence supporting this hypothesis since correlated variability of functionally similar neurons was a major source of spatial inhomogeneities, on a spatial scale that closely matches the anatomical estimates of periodicities in lateral connections and associational input. In contrast, functionally dissimilar neurons showed a strengthening of correlated variability across distance, but did not display any clear periodicity. Interestingly, despite a columnar structure of orientation preference in V1 (Hubel et al. 1978), correlated variability is significantly lower for distant populations, potentially reflecting a much weaker influence of lateral connections. Although a definite answer to the exact relationship between structural and functional connectivity in the PFC could be provided in the future from functional anatomy techniques, the spatial scale consistency across anatomical and functional connectivity measures seems to suggest that indeed, structural connectivity is likely to cluster functionally similar prefrontal populations into local and distant functionally connected ensembles.

The spatial pattern of horizontal connections could be one likely source of the nonmonotonic correlation structure in PFC. Another source could be ascribed to spatially distributed input from associational or thalamic areas to the PFC (35,41,42). Regardless of the underlying mechanism, the impact of spatially clustered, similarly tuned, correlated prefrontal neurons to distant cortical and subcortical targets may facilitate the role of prefrontal cortex in large-scale transmission and integration of information. Specifically, such prefrontal clusters could be thought of as separate channels of information that project to distant cortical and subcortical areas (41,42). Correlated prefrontal output could coordinate these distant targets and therefore contribute to large-scale information processing.

### Spatial structure of prefrontal correlated variability and integrative processing

PFC is a central subnetwork playing a crucial role in cognitive computations due to an increase in the integrative aspect of information processing in higher order cortical areas (43,44). This progressive increase in integrative functions across the cortical hierarchy was recently suggested to be mediated by a similar hierarchy in the timescales of intrinsic fluctuations that arise due to systematic changes in the anatomical structure, like heterogeneous connectivity of local circuitry (6,45,46). A non-monotonic spatial structure of correlated variability differentiates prefrontal functional connectivity from primary sensory areas and could therefore be relevant to the emergence of prefrontal-specific timescales (6,45–47). Network topology was recently suggested to affect timescales since physical distance between connected nodes was shown to increase as time scale lengthened (48).

From a graph-theoretical perspective, a spatially inhomogeneous connectivity profile, combining strong local and long-range functional connectivity, similar to what we observed in the PFC for functionally similar populations, could reflect a network with shorter average path length and higher average clustering coefficient compared to a network with monotonically decreasing correlations and/or uncorrelated long-range functional connectivity (like V1) (49). These topological features are known to facilitate efficient integrative processing (50,51) and could reflect a fundamental characteristic of laterally organized prefrontal microcircuits compared to primary visual cortex where despite positive local correlations, long-range activity on the same spatial scale is uncorrelated (25).

Some recent findings shed light on the spatial functional organization of prefrontal populations that could be critical for integrative processing (52–54). Kiani et al (53) revealed a natural grouping of prefrontal neurons into isolated clusters that remained stable across various conditions (e.g. different epochs of task, spontaneous activity), therefore suggesting that intrinsic lateral connections play a prominent role in shaping functional parcellation in PFC. In another study, Markowitz et al (54) found that different working memory stages are implemented in the PFC by spatially and functionally segregated subnetworks. More importantly, the spiking activity of these subnetworks during working memory is coordinated indicating a distributed network that integrates different aspects of working memory through long range interactions. Our findings, revealing spatially distributed clusters of correlated neurons with similar feature selectivity, provide further evidence for the existence and function of long-range functional interactions within the PFC, which seems to be instrumental for higher-order integrative processing.

### Comparison with previous studies of correlated variability in the prefrontal cortex

Experimental constraints prevented previous studies in dorsolateral areas of the PFC, around the principal sulcus, from capturing a non-monotonic correlation structure (9,55,56). These studies were constrained by a maximum inter-electrode distance of 1 mm, and our findings up to this distance are indeed consistent, showing a decrease in correlations up to 1 mm.

A number of other factors might also have prevented previous studies that used Utah arrays in other areas of the PFC to capture the non-monotonic spatial structure of correlations we report here. Firstly, it is likely that the non-monotonic structure is specific for this particular region of PFC, i.e. vlPFC, since none of these studies involved recordings in the vlPFC but rather in area 8a (i.e. the frontal eye fields) (53,57). The source of this region-specific discrepancy between our results and previous studies (53,57) could be potentially traced to differences in the involvement of various prefrontal regions in visual processing. For example, the probability of finding feature selective neurons (e.g. direction selective neurons) may be higher in the vlPFC compared to area 8a (Hussar and Pasternak, 2009). Since our data validated the spatially nonmonotonic correlation structure during visual stimulation with movie clips and direction of motion, the lack of a similar spatial structure in the frontal eye fields could be due to its differential functional role.

Leavitt et al. (2013) recorded using 4x4mm Utah arrays in area 8A and found a monotonically decreasing correlation structure. However, hardware limitations allowed them to record simultaneously from blocks of only 32 electrodes each time, limiting the spatial coverage that would prevent an extensive examination of the potential spatial anisotropy in area 8A. Kiani et al. (2015), using the same electrode arrays, recorded simultaneously from all 96 electrodes and also reported a monotonic decrease of correlations for multi-unit activity (MUA) for distances up to 4mm. However, the length of electrodes was 1.5mm compared to the 1mm length used in our recordings. Therefore, the monotonically decreasing correlations might be due to layer-specific effects as previously reported in primary visual cortex (58,59).

### Comparison with previous studies of correlated variability in primary visual cortex

Rosenbaum et al. (25) recently provided evidence for a non-monotonic correlation structure in superficial layers (L2/3) of primary visual cortex without strong long-range correlations. In particular, they re-analyzed data collected with Utah arrays during anesthesia and, only after removing the effect of latent shared variability, found that nearby neurons were weakly but significantly correlated, neurons at intermediate distances were negatively correlated and distant neurons were uncorrelated (r_sc_ not different from 0).

There are some major differences between this study and our results from prefrontal recordings on the same spatial scale. Firstly, the average correlation coefficient for distant (3-4 mm apart) neurons in these V1 recordings was not different from zero, which implies an absence of correlation rather than weak correlation between distant populations. In striking contrast, the average magnitude of long-range correlations for the same distance in the awake PFC recordings was a) significantly positive and b) comparable to the magnitude of correlations for nearby neurons. This suggests that long-range (3-4 mm) functional connectivity in PFC is stronger and in fact results in significant long-range correlations compared to the primary visual cortex where, despite a weak non-monotonicity, the average correlated variability between distant neurons is not different from zero. The second and more important difference pertains to the conditions under which the non-monotonic structure was detectable. The Rosenbaum et al. (25) results in V1 suggested an underlying non-monotonic functional connectivity that was washed out by the strong modulatory effects of global state fluctuations (e.g. Up and Down states) observed during anesthesia in macaques (12), and in rodents during anesthesia and quite wakefulness (60,61). Specifically, the non-monotonic correlation structure was revealed only after removing the effect of global latent fluctuations via Gaussian Process Factor Analysis (GPFA). This suggests that a non-monotonic structure in the primary visual cortex should be directly detectable in data recorded from awake animals where the anesthesia-induced global fluctuations are absent. However, to the best of our knowledge, until now there is no direct experimental evidence in awake V1 recordings. In contrast, Ecker et al. (24) found a flat correlation structure in awake V1 recordings using tetrode arrays, which was also revealed after removing latent fluctuations from anesthetized recordings using the same technique (12). Regardless of the underlying reason for this discrepancy (e.g. layer specificity or the effect or number of samples), our recordings in the PFC provide the first direct evidence for a non-monotonic, long-range correlation structure during wakefulness, without the need for removing latent sources of covariance, i.e. without application of GPFA or any other similar tool involving theoretical assumptions like stationarity of responses or the number of latent factors that contribute in driving correlated variability.

## Conclusion

Overall, our results suggest that the mesoscopic functional connectivity architecture of vlPFC is fundamentally different compared to early sensory cortices such as V1 or V4. Correlated variability in the vlPFC is spatially non-monotonic and a major source of non-monotonicity is the spatial pattern of correlations between neurons with similar functional properties. A non-monotonic functional connectivity profile with strong and equivalent local and long-range interactions might reflect the underlying machinery for large-scale coordination of distributed information processing in the prefrontal cortex.

## Experimental Procedures

### Electrophysiological recordings

Extracellular electrophysiological recordings were performed in the inferior convexity of the lateral PFC of 2 anesthetized and 2 awake adult, male rhesus macaques (*Macaca mulatta*) using Utah microelectrode arrays (Blackrock Microsystems, Salt Lake City, Utah USA; (37). The array (4x4mm with a 10 by 10 electrode configuration and inter-electrode distance of 400?m) was placed 1 - 2 millimeters anterior to the bank of the arcuate sulcus and below the ventral bank of the principal sulcus, thus covering a large part of the inferior convexity in the ventrolateral PFC (Figure 1A). For the awake experiments, monkeys were implanted with form-specific titanium head posts on the cranium after modelling the skull based on an anatomical MRI scan acquired in a vertical 7T scanner with a 60cm diameter bore (Biospec 47/40c; Bruker Medical, Ettlingen, Germany). The methods for surgical preparation and anesthesia have been described in previous studies (62–64). All experiments were approved by the local authorities (Regierungspräsidium) and were in full compliance with the guidelines of the European Community (EUVD 86/609/EEC) for the care and use of laboratory animals.

### Data acquisition and spike sorting

Broadband neural signals (0.1-32 kHz in the anesthetized recordings, and 0.1-30 kHz in the awake recordings) were recorded using a Neuralynx (Digital Lynx) data acquisition system for the anesthetized recordings and Neural Signal Processors (Blackrock Microsystems) for the awake recordings.

In the anesthetized data, to detect spiking activity we first band–pass filtered (0.6-5.8 kHz) the broadband raw signal using a minimum–order finite impulse response (FIR) filter (65) with 65dB attenuation in the stop–bands and less than 0.002dB ripple within the pass–band. A gaussian distribution was fit to randomly selected chunks of the filtered signal to compute the noise variance, and the amplitude threshold for spike detection was set to 5 times the computed variance. Spike events with inter–spike intervals less than a refractory period of 0.5ms were eliminated. Those events that satisfied the threshold and refractory period criteria were kept for spike sorting.

In the awake experiments, broadband data were filtered between 0.3-3 kHz using a 2^nd^ order Butterworth filter. The amplitude for spike detection was set to 5 times the median absolute deviation (66). The criterion for rejection of spikes was the same as described above. All the collected spikes were aligned to the minimum. For spike sorting, 1.5 ms around the peak, i.e. 45 samples were extracted.

Automatic clustering to detect putative single neurons in both the awake and anesthetized data was achieved by a Split and Merge Expectation-Maximisation (SMEM) algorithm that fits a mixture of Gaussians to the spike feature data which consisted of the first three principal components. For the anesthetized data, the SMEM algorithm by Ueda et al was used (67). Details of the spike sorting method used in this study have been described in other papers using tetrodes (24,68). For the awake data, the KlustaKwik algorithm (69,70) was employed. The spike sorting procedure was finalized in both cases through visual inspection using the program Klusters (71).

### Visual stimulation

In anesthetized recordings, full-field visual stimulation of 640 × 480 resolution with 24-bit true colour at 60 Hz for each eye was presented using a Windows machine equipped with an OpenGL graphics card (Wildcat series; 3DLABS). We used 10 second epochs from a commercially available movie (Star Wars Episode 1, the Battle of Naboo). Hardware double buffering was used to provide smooth animation. The experimenter’s monitor and the video interface of a fiber-optic stimulus presentation system (Silent Vision; Avotec) were driven by the VGA outputs. The field of view was 30 (horizontal) × 23 (vertical) degrees of visual angle, and the focus was fixed at 2 diopters. Binocular presentation was possible through two independently positioned plastic, fiber-optic glasses; however in this study we used monocular stimulation (either left or right eye). The contact lenses for the eyes had matched diopter with an Avotec projector, to focus images on the retina. Positioning was aided by a modified fundus camera (RC250; Carl Zeiss) with an attachment to hold the projector on the same axis of the camera lens. After observing the foveal region, the projector was fixed relative to the animal.

In the awake recordings, the visual stimuli were generated by in-house software written in C/Tcl and used OpenGL implementation. Stimuli were displayed using a dedicated graphics workstation (TDZ 2000; Intergraph Systems, Huntsville, AL, USA) with a resolution of 1,280 × 1,024 and a 60Hz refresh rate. An industrial PC with one Pentium CPU (Advantech) running the QNX real-time operating system (QNX Software Systems) controlled the timing of stimulus presentation, digital pulses to the Neuralynx system (anesthetized) or the Blackrock system (awake), and acquisition of images. Eye movements were captured using an IR camera at 1kHz sampling rate using the software iView (SensoriMotoric Instruments GmBH, Germany). They were monitored online and stored for offline analysis using both the QNX-based acquisition system and the Blackrock data acquisition system. In the anesthetized recordings, neural activity was recorded in 200 trials of repeated stimulus presentation. Each trial consisted of the same 10s long movie clip, followed by 10s of a blank screen (intertrial). In the awake experiments, two monkeys were trained to fixate on a red square of size 0.2° of visual angle subtended on the eye about 45cm from the monitors, and maintain fixation within a window of 1.5-2° of visual angle. The location of the red fixation square was adjusted to the single eye vergence of each individual monkey. After 300ms of fixation, a moving grating of size 8°, moving at a speed of 12° (monkey H) and 13° (monkey A) per second with a spatial frequency of 0.5 cycles/degree of visual angle and at 100% contrast was presented for 1000ms. The gratings encompassed 8 different directions of motion viz. 0°, 45°, 90°, 135°, 180°, 225°, 270° and 315° (Figure 1B), pseudo-randomized within a block of 8 trials. After 1000ms, a 300ms stimulus-off period preceded the completion of the trial. The monkeys were given a liquid reward (either water or juice) at the end of the trial, if they maintained fixation within the specified fixation window during the entire duration of the trial. Every successful trial was followed by a 1000ms inter-trial period. On average, we found 32 +/-5% of all recorded neurons to be visually modulated. The stimuli, although presented through a stereoscope (due to the data being collected on the same day with other experiments requiring dichoptic viewing conditions), were always presented monocularly in the left eye to remain consistent with the monocular stimulation protocol used in the anesthetized recordings. In both anesthetized and awake recordings, to ensure accurate control of stimulus presentation, a photodiode was attached to the experimenter’s monitor permitting the recording of the exact presentation time of every single frame.

In the awake recordings, spontaneous activity datasets were collected on days different from those of the task recordings. The monkeys were allowed to move their eyes freely, or have their eyes closed. The recording chamber was sound-resistant and dark. In the anesthetized recordings, spontaneous activity datasets were recorded between periods of visual stimulation. In both the awake and anesthetized recordings of spontaneous activity, the monitors were turned off. The duration of each spontaneous activity dataset was between 40 to 80 minutes.

### Tuning functions and signal correlations

Tuning curves for each detected single unit were computed by averaging the firing rate across trials for each of the 8 presented directions of motion. Signal correlations, defined as the correlation coefficient between the tuning curves of a neuronal pair, were also computed (7). In addition to classical tuning curves (direction and orientation selectivity), other types of tunings such as inverted tunings, for example, have also been reported in the electrophysiological studies of the macaque prefrontal cortex (72). Because of this variability in the observed tuning properties of detected single units, signal correlation provides a more general measure of response similarity and therefore it was used to investigate the correlation structure that arises from this functional similarity.

### Spike count correlations

To compute spike count correlation (*r*_*sc*_) during the anesthetized state, we divided the period of visual stimulation into 10 periods, each being 1000ms long, and considered these periods as different successive stimuli. The intertrial period was also binned in the same way. In the awake data, visual stimulation and intertrial periods were 1000ms long each, thus being consistent with the anesthetized experiments. We estimated spike counts over 1000ms due to the stimulus length used in previous studies of correlated variability. In spontaneous datasets (both anesthetized and awake), the entire length of the recording epoch was split into periods of 1000ms that were treated as a trial.

The spike count correlation coefficients were computed similar to previous studies in primary visual areas (10,24,62). First, for each condition (either presentation of each moving grating in awake experiment or a single bin of movie clip in the anesthetized experiment), we normalized the spike counts across all trials by converting them into *z*-scores (10). For each pair, we computed the Pearson’s correlation coefficient for the two vectors *z*_*i*_ and *Z*_*j*_ as following:

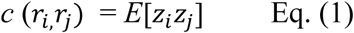

After computing *c*(*r*_*i*_, *r*_*j*_ for each condition, we averaged across conditions to obtain the correlation value. Equivalently, one can concatenate z-scores for all the conditions in long vectors and find the expectation of their product. To account for possible non-physiological correlations between detected neurons, which could happen for example, due to shorts between recording electrodes, a threshold of 5 standard deviations above the mean correlation value was set and the outliers were discarded.

### Quantification of spatial inhomogeneities in correlated variability

We quantified the inhomogeneity in the spatial structure of correlated variability across different conditions and states by computing the mean of the absolute rate (i.e. first differential) of correlation changes across lateral distance. To estimate the first differential with respect to distance, we subtracted the mean correlation values of consecutive bins that were significantly different (Wilcoxon rank-sum test, alpha level 0.05). If no significant change between two consecutive bins was observed, the derivative at that point was set to zero.

### Curve fitting procedures

A two parameter line (*y = ax + b*) and a three parameter quadratic function (*y = ax^2^ + bx + c*) were fit via a minimization of the least squared error to the results in Figure 6B and 6D using the in-built Curve Fitting Toolbox in MATLAB 2016b. The chosen functions (39) were fit to the mean noise correlation functions which were weighted by the standard error of the mean of each data point as the individual data points spanned varying number of observations.

## Acknowledgments

This study was supported by the Max Planck Society. We thank Britni Crocker and Zeynab Razzaghpanah for help with pre-processing of the data and spike sorting, and Yusuke Murayama and the other technical and animal care staff for excellent technical assistance. We also thank Prof. Andreas Tolias for help with the initial implantations of the Utah arrays and Dr. Michel Besserve and Christos Constantinidis and Rodrigo Quian Quiroga for their comments on a previous version of this manuscript.

## Author Contributions

Conceptualization, T.I.P.; Methodology, S.S., A.D., V.K. and T.I.P.; Software, S.S., A.D., T.I.P. and J.W.; Formal Analysis, S.S., A.D. and T.I.P.; Investigation, V.K., A.D., T.I.P., S.S. and N.G.H.; Resources, N.K.L.; Data Curation, A.D., T.I.P., V.K., and S.S.; Writing – Original Draft, T.I.P., S.S., and A.D.; Writing – Review & Editing: V.K., A.D., T.I.P., N.G.H. and N.K.L.; Visualization, S.S., A.D, V.K. and T.I.P.; Supervision and Project administration, T.I.P.; Funding acquisition, N.K.L.

---

**Table S1: Related to figure 2.** Summary of p-values in the anesthetized state for comparison of correlated variability in three key distance bins mentioned in the main text, viz 0.5mm, 2.5mm and 3.5mm

| Distance bins | Bonferroni adjusted alpha | Visual stimulation all pairs (figure 2B red) |
| --- | --- | --- |
| 0.5 mm vs 2.5 mm | 0.0167 | $< 10^{-10}$ |
| 2.5 mm vs 3.5 mm | 0.0167 | 0.004 |
| 0.5 mm vs 3.5 mm | 0.0167 | 0.0008 |

**Table S2: Related to figure 2.** Summary of p-values in the awake state for comparison of correlated variability in three key distance bins mentioned in the main text. (1mm in during visual stimulation, 0.5mm during intertrial and spontaneous states)

| Distance bins | Bonferroni adjusted alpha | Visual stimulation all pairs (figure 2B red) | Intertrial all pairs (figure 2B black) | spontaneous all pairs (figure 2B dark blue) |
| --- | --- | --- | --- | --- |
| 0.5 or 1.0 mm vs 2.5 mm | 0.0167 | 0.0038 | 0.3 | 0.6 |
| 2.5 mm vs 4.0 mm | 0.0167 | 0.0079 | 0.25 | $> 0.1$ |
| 0.5 or 1.0 mm vs 4.0 mm | 0.0167 | $> 0.3$ | 0.8 | 0.33 |

**Table S3: Related to figure 6.** Summary of p-values in the awake state among functionally similar and dissimilar populations for comparison of correlated variability in three key distance bins mentioned in the main text. (0.5mm, 2.5mm and 4mm)

| Distance bins | Visual stimulation<br>pairs with $r_{\text{signal}} > 0.1$<br>(figure 6B green) | Visual stimulation<br>pairs with $r_{\text{signal}} < 0.1$<br>(figure 6B blue) | Intertrial<br>pairs with $r_{\text{signal}} > 0.1$<br>(figure 6D green) |
| --- | --- | --- | --- |
| 0.5mm vs 2.5mm | 0.07 | 0.9 | 0.3 |
| 2.5mm vs 4mm | 0.1 | 0.24 | 0.9 |
| 0.5mm vs 4mm | 0.08 | 0.4 | 0.6 |

## Supplemental Experimental Procedures

Surgical preparation and anesthesia

In the anaesthetized experiment, the animals were initially premedicated with glycopyrrolate (0.01 milligrams (mg) per kilogram of body weight (kg), i.m.) and ketamine (15 mg per kg, i.m.). Next, an intravenous catheter was inserted and vital monitors (HP OmniCare/CMS; Hewlett Packard; electrocardiogram, noninvasive blood pressure, CO2, SpO2, temperature) were connected. The monkeys were preoxygenated and anesthesia was induced with fentanyl (3 micrograms (?g) per kg), thiopental (5 mg per kg), and succinylcholine chloride (3 mg per kg) for the intubation of the trachea. The animals were ventilated using an Ohmeda anesthesia machine (Ohmeda), maintaining an end-tidal CO2 of 33 mmHg and oxygen saturation over 95%. Balanced anesthesia was maintained with remifentanil (typical, 1 ?g per kg per minute). Mivacurium (5 mg per kg per hour) was used for muscle relaxation. Body temperature was kept constant, and lactated Ringer’s solution was given at a rate of 10 milliliters (ml) per kg per hour. During the entire experiment, the vital signs of the monkey and the depth of anesthesia were continuously monitored.

Drops of 1% ophthalmic solution of anticholinergic cyclopentolate hydrochloride were given to each eye to prevent accommodation of the lens and dilation of the pupil. Refractive errors were measured and contact lenses (hard PMMA lenses; Wöhlk) were put on the monkey’s eyes with continuous drops of saline throughout the experiment to prevent the eyes from drying. The lenses with the appropriate dioptric power were used to bring the animal’s eyes into focus on the stimulus plane.

**Figure S1: Related to Figure 2:**
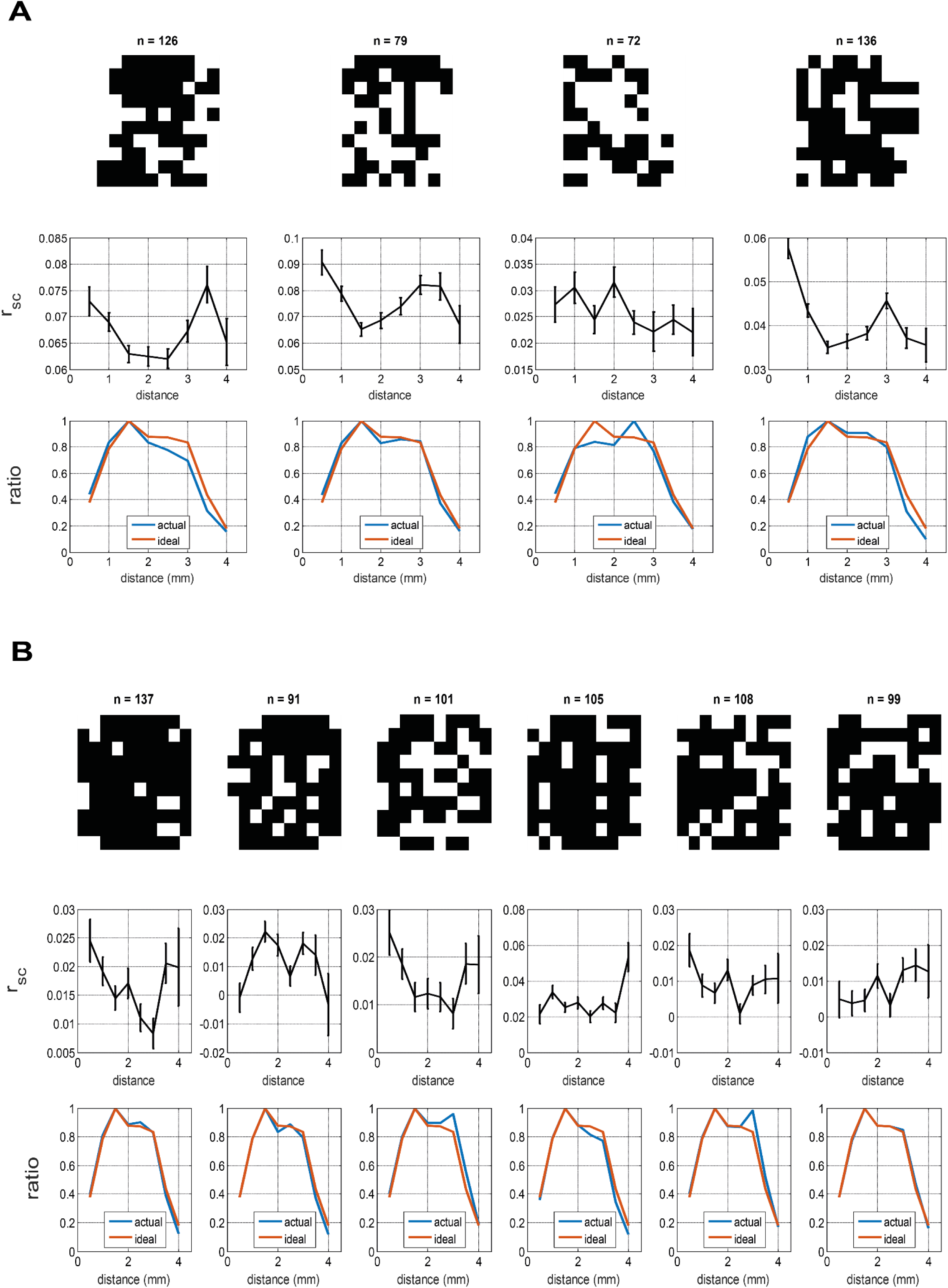
Spatial distribution of single units in anesthetized and awake state recordings. (A) The first row depicts the spatial coverage of the detected single units (*n*) on the Utah array (10 x 10 array of electrodes) for four individual datasets from the anesthetized recordings (each column corresponds to an individual dataset). Black pixels indicate channels where at least one single unit was detected. In the second row, similar to Figure 2A-B in the main text, correlated variability is depicted as a function of distance during visual stimulation for the same datasets. In the third row we plot the spatial distributions of the recorded pairs as a function of distance on the Utah arrays (*actual*) for individual datasets. In order to test if the spatial structure of spike count correlations was an artefactual reflection of the spatial distribution of isolated neurons in our recordings, we compared the actual distribution of the pairs with an “ideal” distribution across distance bins. This ideal distribution is derived from a simulated dataset where we could detect equal number of neurons on all sites, indicating the best possible sampling one can achieve with the Utah arrays. To simulate this ideal distribution, we consider a maximum number of detected single neurons on all sites and find the distribution of pairs (aka *ideal* distribution). The largest number of single units among all sites in the recorded dataset is used as the maximum number. For example, if in a given dataset, *n* single units were the largest number of single units across all electrodes, we consider a distribution derived from *n* single units across all sites. The choice of this maximum is not crucial as distributions are normalized. Red and orange traces indicate the actual and ideal distribution of number of pairs (normalized to the peak) in four datasets across all distance bins. The spatial spread of the recorded single units is not significantly different from an ideal distribution. (B) Similar to figure (A), but for awake state datasets.

**Figure S2: Related to Figure 2:**
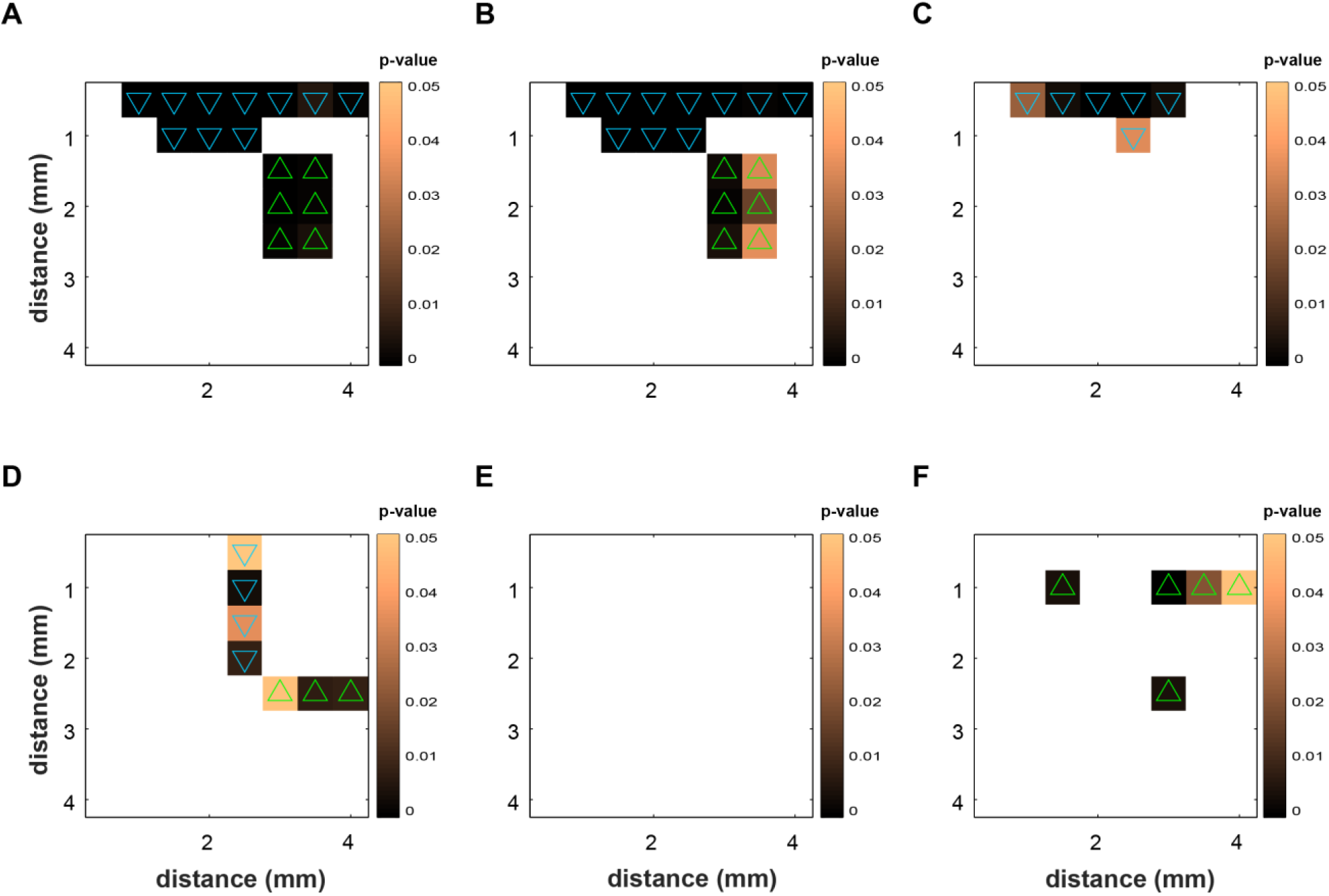
Pairwise comparison of correlations across distance bins. (A-F) Pairwise comparison of correlation distributions (Wilcoxon rank-sum test) across all distance bins for different states and conditions: anesthetized - visual stimulation (A), intertrial (B), spontaneous activity (C) and awake - visual stimulation (D), intertrial (E), spontaneous activity (F). Each plot is an 8 x 8 symmetric matrix (therefore only the upper triangular part is kept) corresponding to all possible pairwise comparisons of 8 distance bins. For example, the p-value of the comparison between 0.5 and 2.5 mm is encoded in the colour of the pixel located in row 1 and column 5 of the matrix. All coloured pixels indicate a significant change since pixels without significant differences (p>0.05) were removed (white pixels). Green triangles indicate a significant increase in the correlation values, while blue triangles denote a significant decrease.

**Figure S3: Related to Figure 2:**
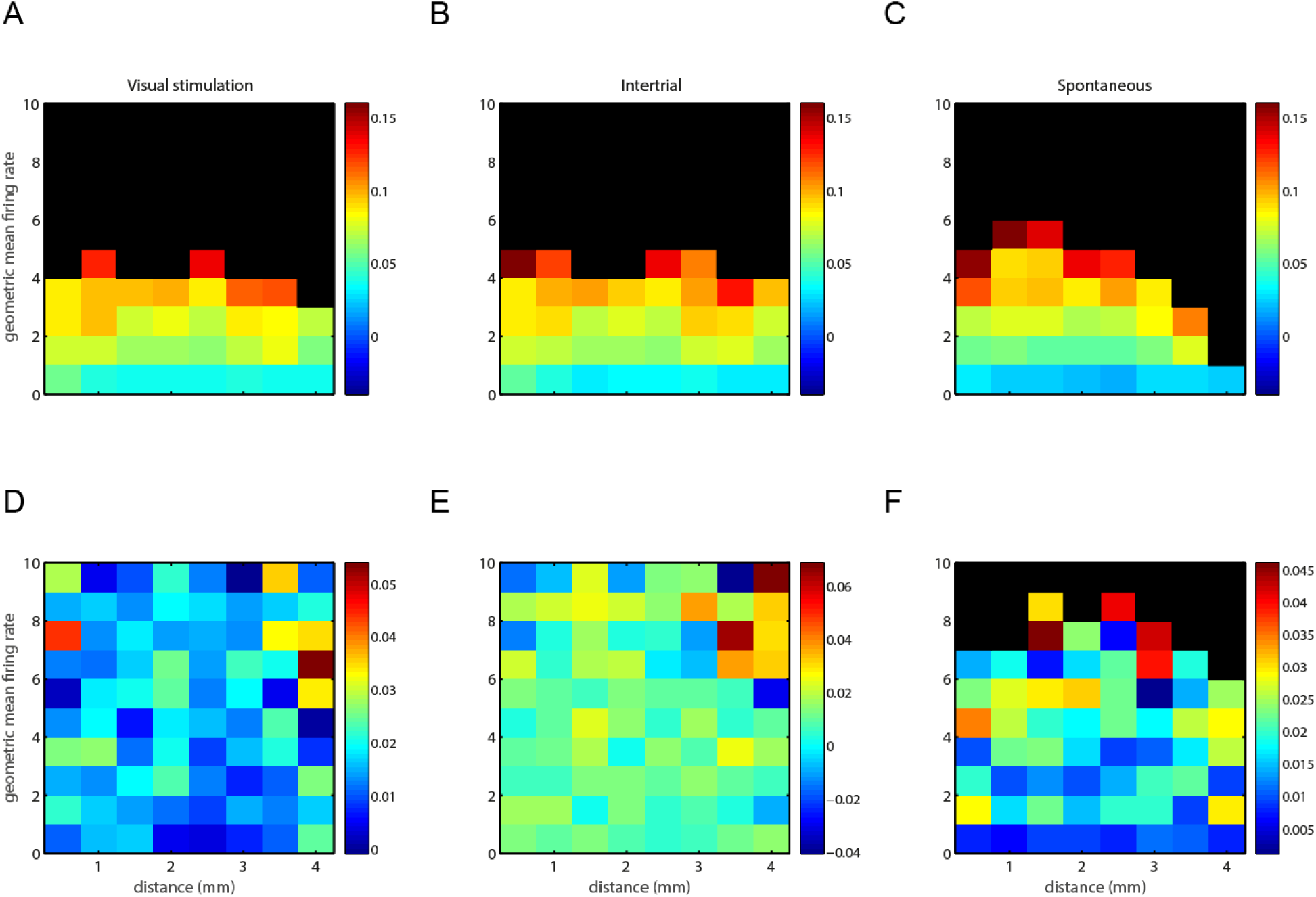
Correlated variability as a function of distance for populations with matched geometric mean firing rates. To control for spatial variations in firing rates giving rise to the non-monotonic structure of correlated variability, we plot correlated variability as a function of distance and geometric mean of the firing rates during different states and conditions: anesthetized - visual stimulation (A), intertrial (B), spontaneous activity (C) and awake - visual stimulation (D), intertrial (E), spontaneous activity (F). The colour of each pixel indicates the average correlated variability for pairs whose geometric mean firing rate and distance landed in the specific bin. Therefore, each row contains pairs with similar firing rate geometric mean. Pixels containing less than 10 pairs are removed (black pixels). Correlated variability values are indicated by the colour bar at the right of the panel. Data were not smoothed.

**Figure S4: Related to Figure 2:**
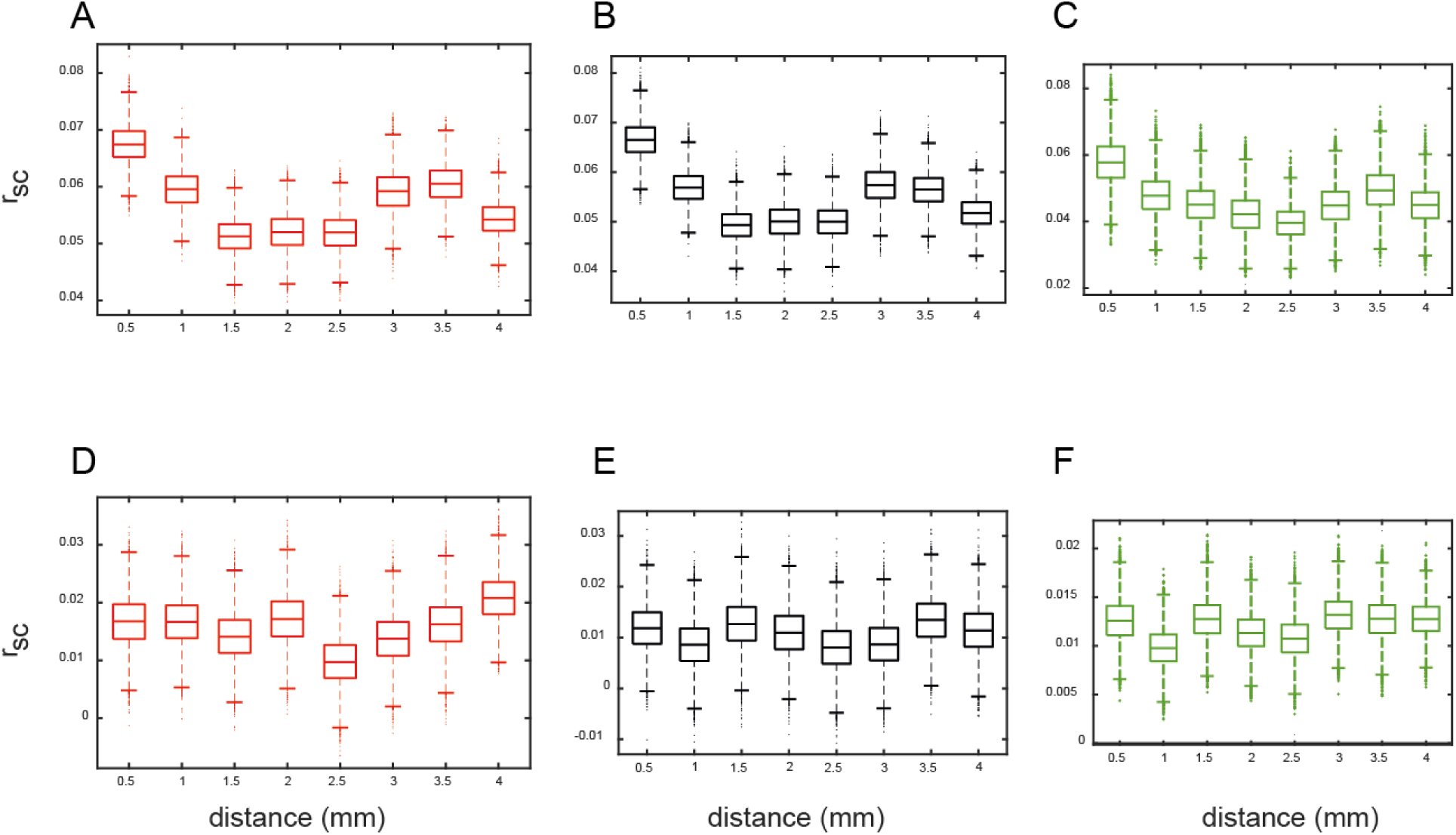
Bootstrapping for the spatial sampling. To control for whether the intrinsic non-uniformity of spatial sampling with Utah array influences the nonmonotonic structure of correlated variability, we used a Bootstrap analysis for different states and conditions: anesthetized - visual stimulation (A), intertrial (B), spontaneous activity (C) and awake - visual stimulation (D), intertrial (E), spontaneous activity (F). Each figure depicts the box-plot of the mean of the thus derived pseudo-datasets across all distance bins. For each state-condition (A-F) we bootstrapped 10000 times. We obtained a set of G (=10000) draws from the distribution of all pairs belong to a particular distance bin. For each draw, we constructed a pseudo-dataset where the size of each pseudo-dataset is limited by the sample size of the distance bin that contains the least number of pairs. Therefore all distance bins contain an equal number of samples in any draw (usually the largest distance bin i.e. 3.75mm-4.25mm). This analysis suggests that the equalized resampling of the pairs across distance bins show similar results (figure 2). Therefore, the non-monotonic structure in correlated variability could not be ascribed to the non-uniformity of the spatial sampling of the Utah arrays.

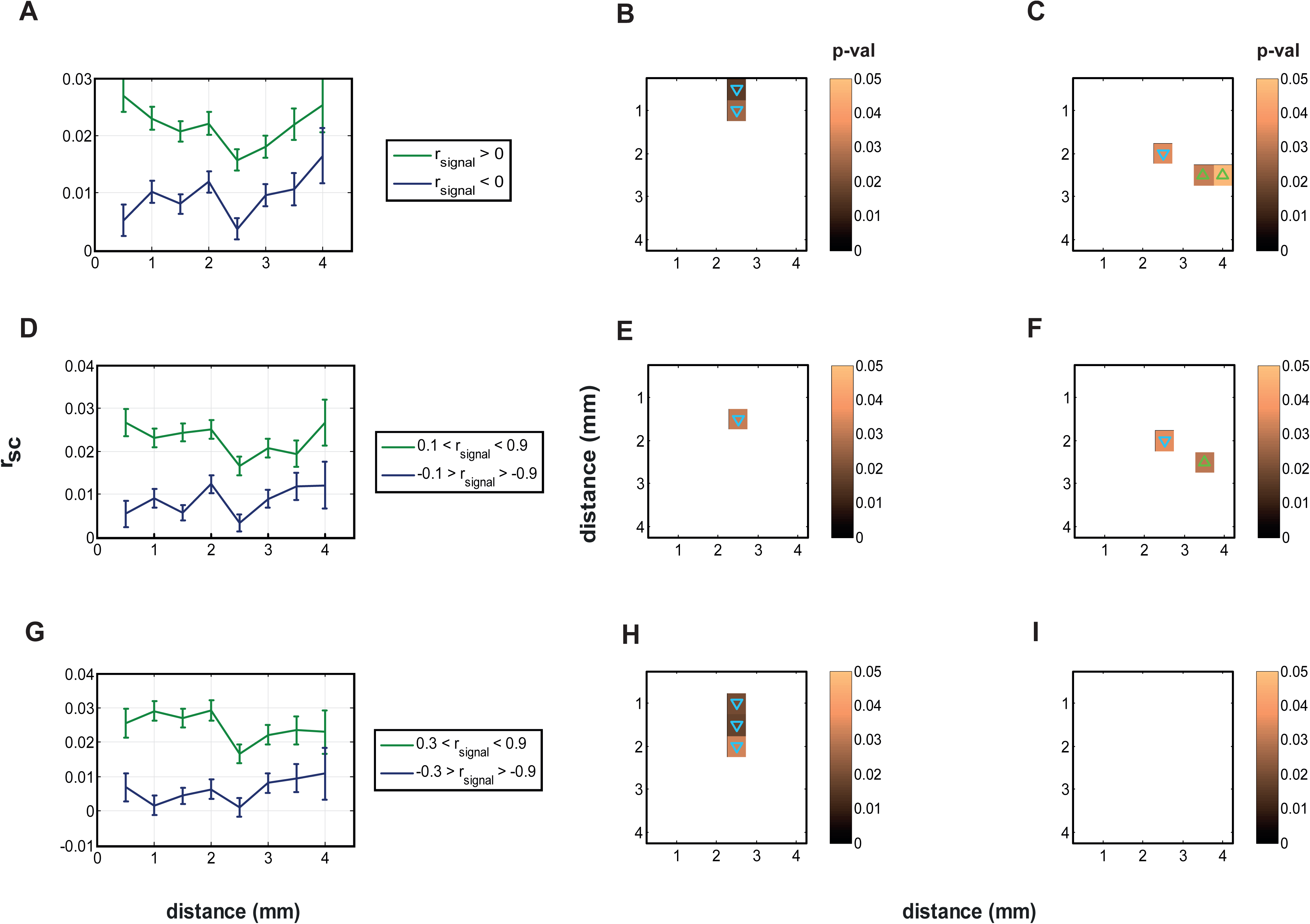

